# Discovery of new mitochondrial DNA segregation machinery components in *Trypanosoma brucei*: a comparison of three approaches

**DOI:** 10.1101/2019.12.28.889931

**Authors:** Hélène Clémentine Margareta Baudouin, Laura Pfeiffer, Torsten Ochsenreiter

## Abstract

*Trypanosoma brucei* is a single celled eukaryotic parasite and the causative agent of human African sleeping sickness and Nagana in cattle. Aside from its medical relevance *T. brucei* has also been key to the discovery of several general biological principles including GPI-anchoring, RNA-editing and trans-splicing. The parasite contains a single mitochondrial organelle with a singular genome. Recent studies have identified several molecular components of the mitochondrial genome segregation machinery (tripartite attachment complex, TAC), which connects the basal body of the flagellum to the mitochondrial DNA of *T. brucei*. The TAC component in closest proximity to the mitochondrial DNA is TAC102. Here we apply and compare three different approaches (proximity labeling, Immunoprecipitation and yeast two-hybrid) to identify novel interactors of TAC102 and subsequently verify their localisation. Furthermore, we establish the direct interaction of TAC102 and p166 in the unilateral filaments of the TAC.

**Subject area:** biochemistry, molecular biology, cellular biology

## Introduction

*Trypanosoma brucei*, a protozoan parasite, is the causative agent of human African sleeping sickness and Nagana in cattle (Achcar, Kerkhoven, and Barrett 2014; Büscher et al. 2017). The single celled eukaryote belongs to group of Kinetoplastea, that is characterised by the presence of a singular yet complex mitochondrial genome consisting of a DNA network referred to as kinetoplast DNA or kDNA (Adl et al. 2005; Shapiro and Englund 1995; Vickerman 1977). The kDNA in *T. brucei* consists of 25 almost identical maxicircles (23 kbp) that are linked to thousands of 1 kbp size minicircles, which are in turn catenated to each other. The maxicircles encode for 18 protein genes that are involved in oxidative phosphorylation and the mitochondrial ribosome as well as two ribosomal RNAs. Most (12) of the maxicircle protein coding genes are pseudogenes and require posttranscriptional insertion and/or deletion of uridine residues in order to be translatable (Hajduk and Ochsenreiter 2010; Benne et al. 1986). This process, called RNA editing, requires a 34S ribonucleotide protein complex consisting of more than 20 different proteins as well as small, 50-70 nucleotide, guide RNAs (gRNAs) that define the editing pattern (Stuart et al. 2005; Read, Lukeš, and Hashimi 2016; Blum, Bakalara, and Simpson 1990). The gRNAs are encoded on the minicircles of the network. A recent study showed that the *T. brucei* mitochondrial genome harbors about 400 different minicircle sequences in the network coding for 1300 gRNA genes (Cooper et al. 2019).

The complexity of replicating the kDNA rivals its structure, and although more than 30 components have been characterised, the compendium of the kDNA replication machinery is far from complete (Jensen and Englund 2012; Klingbeil 2004). Since the parasite contains a single mitochondrion with one genome per cell, proper segregation of these two entities is critical for cell proliferation. In the bloodstream form parasite the mitochondrion grows in two regions anterior and posterior of the nucleus building a network that is pruned prior to separation in the two daughter cells (Jakob et al. 2016). The segregation of the replicated kDNA network is carried out by the tripartite attachment complex (TAC) that is composed of three parts; (i) the exclusion zone filaments, connecting the basal body of the flagellum to the outer mitochondrial membrane, (ii) the two differentiated mitochondrial membranes and (iii) the unilateral filaments connecting the inner mitochondrial membrane to the kDNA (Ogbadoyi 2003). The current model of the TAC contains 13 components (Schneider and Ochsenreiter 2018) four of which (p197, BBA4, Mab22 and TAC65) are localised to the exclusion zone filaments (Gheiratmand et al. 2013; Bonhivers et al. 2008; Käser et al. 2016). Six components are in the differentiated mitochondrial membranes, four in the outer mitochondrial membrane TAC60, TAC42, TAC40 and pATOM36 (Schnarwiler et al. 2014; Käser et al. 2016, 2017) and two components are likely associated with the inner mitochondrial membrane (p166, AEP1, (Zhao et al. 2008; Ochsenreiter and Hajduk 2006)). TAC102 is a protein of the unilateral filaments and the most proximal component to the kDNA (Trikin et al. 2016; Hoffmann, Jakob, and Ochsenreiter 2016). A number of additional components that display multiple localisations including in the TAC, like the E2 subunit of the α-ketoglutarate dehydrogenase and the tubulin-binding cofactor C protein, have been identified (Sykes and Hajduk 2013; André et al. 2013). The assembly of the TAC occurs *de novo* from the base of the flagellum towards the kDNA in a hierarchical manner such that kDNA proximal components depend on the proper assembly of the kDNA distal components (Hoffmann et al. 2018; Schneider and Ochsenreiter 2018). In order to identify novel interactors of TAC102, we used three different approaches with TAC102 as the bait: (i) proximity-dependent biotin identification (BioID), (ii) immunoprecipitation with a monoclonal TAC102 antibody and (iii) yeast two-hybrid. BioID was first used for the identification of protein-protein interactions in mammalian cells (Roux et al. 2012). It involves the expression of a protein of interest fused to a modified version of a bacterial biotin ligase (BirA*, (Kwon and Beckett 2000)), in order to identify protein partners in close proximity. The enzyme adds an ATP to biotin in order to form biotinoyl-5’-AMP, which is highly reactive (Lane et al. 1964) and leads to biotinylation of all proteins in a radius of 20 nm around BirA*. The biotinylated proteins can then be purified using streptavidin beads and are identified by mass spectrometry. The BioID approach was previously applied to identify interacting partners of the bilobe protein TbMORN1 a key component of the cytoskeleton structure close to the flagellar pocket of *T. brucei* (Morriswood et al. 2013). Immunoprecipitation has widely been used to characterise protein-protein interactions in *T. brucei*. However, in most studies the protein of interest has been tagged, which can potentially lead to a change in protein expression levels as well as interference of the tag with the protein’s function. In this study we used a monoclonal antibody raised against a region in the C-terminus of TAC102. This antibody was previously shown to be highly specific for TAC102 in fixed cells and native/denatured protein extracts (Trikin et al. 2016; Hoffmann et al. 2018). The third approach we applied was a yeast two-hybrid screens that relies on the reconstitution of a functional transcription factor when two peptides of interest interact (Fields and Song 1989). One of the advantages of the yeast two-hybrid screen is its ability to identify interaction domains through the expression of parts of the bait/prey proteins. However for the same reason false positive/negative interactions are possible since the peptides might fold differently than the entire protein. Yeast two-hybrid screens have successfully been used in *T. bruce*i, albeit much less frequently than immunoprecipitation approaches. Two examples reporting protein-protein interactions in the parasite are the characterisation of a sumoylation factor interacting with the transcription machinery regulating VSG expression and the description of mitochondrial protein-protein interactions in the RNA editing accessory complex MRB1 (López-Farfán et al. 2014; Ammerman et al. 2012).

Here we compare the three approaches for the discovery of novel interactors of TAC102, an essential component of the mitochondrial DNA segregation machinery in *T. brucei*. We identify nine proteins to be enriched in both the BioID and the immunoprecipitation approach, three of which we localize to the TAC/kDNA region. The yeast two hybrid and the immunoprecipitation approach identified p166 a previously described TAC component as the main interactor of TAC102.

## Results

### Myc-BirA*-TAC102 is active and colocalises with the endogenous TAC102

The Myc-BirA*-TAC102 fusion construct was integrated into the ribosomal array in PCF cells allowing for inducible expression through the addition of tetracycline. After six hours of induction we visualised the expression of the Myc-BirA*-TAC102 fusion protein by immunofluorescence microscopy (Figure 1A) and found it to be colocalized with the endogenous TAC102 (Tb927.7.2390) in the posterior region of the mitochondrion close to the kDNA. Furthermore, based on visual inspection of the micrographs TAC102 and the Myc-BirA*-TAC102 fusion protein were expressed at comparable levels after six hours of induction (Figure 1A). In order to test if the fusion protein associates with the TAC structure similarly as we previously showed for the endogenous TAC102 (Trikin et al. 2016), we evaluated the solubility using increasing amounts of the non-ionic detergent digitonin during a biochemical fractionation. For this analysis the Myc-BirA*-TAC102 fusion protein was expressed for 24 hours. The supernatants from the digitonin fractions were separated on a SDS-PAGE for western blot analysis (Figure 1B) demonstrating that the majority of Myc-BirA*-TAC102 are solubilized at similar concentration of digitonin (0.2%) as previously described for the endogenous TAC102 (Trikin et al. 2016). Aside from the signal for the Myc-BirA*-TAC102 fusion protein (about 135 kDa) and the weaker signal for TAC102 (102 kDa) we also detected several additional signals (see discussion).

**Figure 1.**
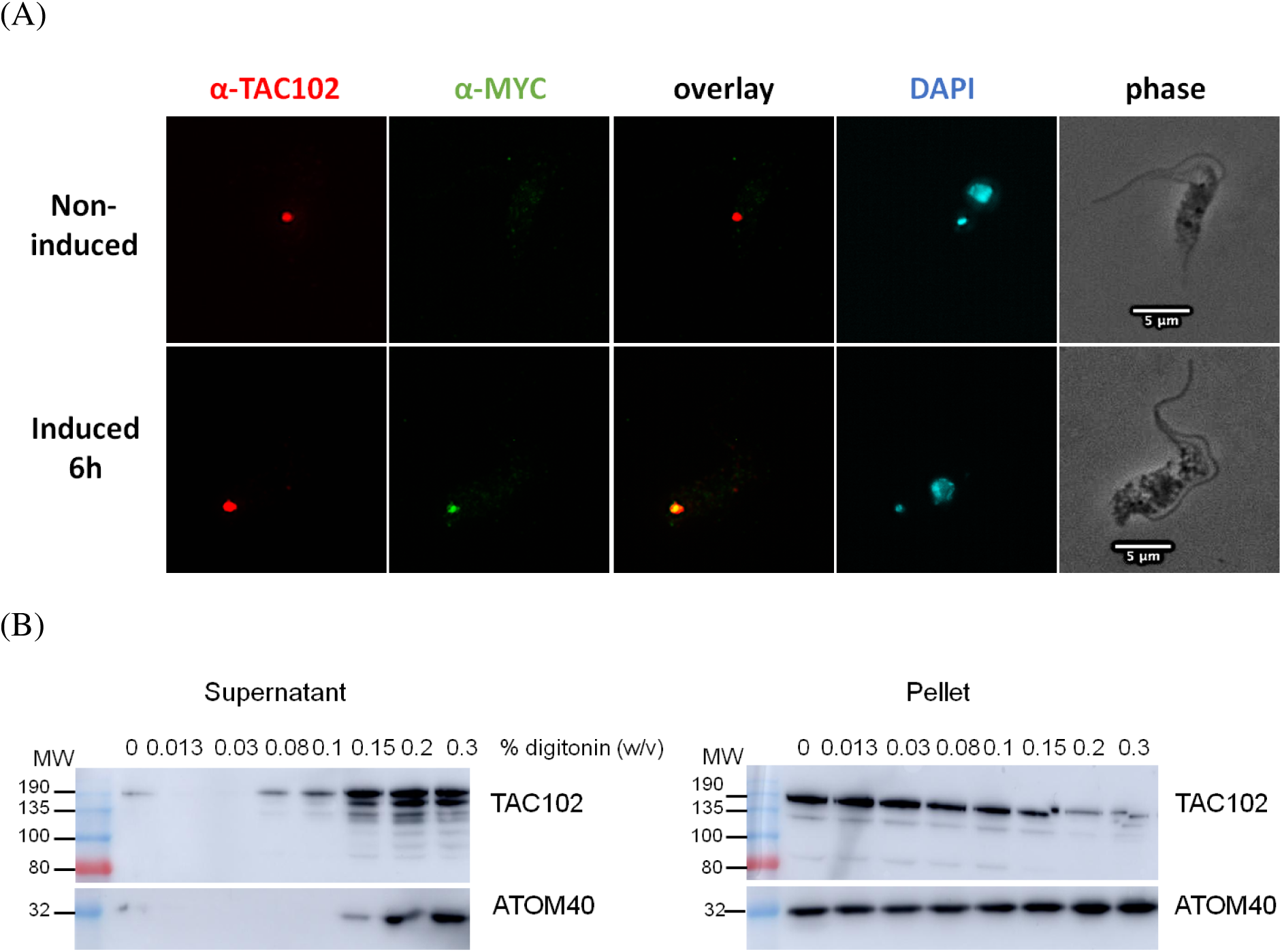
Characterisation of the Myc-BirA*-TAC102 cell line. (A) Immunofluorescence microscopy pictures of Myc-BirA*-TAC102 PCF cells. A monoclonal antibody is used to visualise TAC102 (red). An anti-Myc antibody is used to visualise Myc-BirA*-TAC102 (green). DAPI stains nuclei and kDNA (blue). (B) Western blot analysis of digitonin fractionated cell extracts from the Myc-BirA*-TAC102 cell line (PCF) after 24 hours of induction with tetracycline. Myc-BirA*-TAC102 and the endogenous TAC102 are detected using the monoclonal anti-TAC102 antibody. Molecular weights (MW) are in kDa. As control we probed for the mitochondrial membrane protein ATOM40.

### Myc-BirA*-TAC102 is able to biotinylate proteins

In order to test the activity of the Myc-BirA*-TAC102 fusion protein inside the mitochondrial organelle, we induced its expression in procyclic form cells and then evaluated the biotinylation pattern of the total cell extract by western blot (Figure 2A). After induction of the fusion protein, we could detect an increase of biotinylated proteins when compared to the non-induced control. The major band visible between 135 and 190 kDa is likely the auto-biotinylated Myc-BirA*-TAC102 fusion protein (Figure 2A). The overall level of biotinylation further increased when exogenous biotin was added to a final concentration of 50 µM. Most biotinylated proteins were soluble in the lysis buffer (Fraction S1, Figure S1C). While they are readily detectable by western blot the overall amount of biotinylated proteins is low as seen on Coomassie stained polyacrylamide gels (Fraction B1, Figure S1B). The localisation of the biotinylated proteins was analysed by epifluorescence microscopy using a streptavidin-conjugated fluorophore (Alexa Fluor 488, Figure 2B). For this the cells were first incubated with biotin for 24 hours before adding tetracycline to induce expression of Myc-BirA*-TAC102 for five hours. The streptavidin-conjugated antibody recognised proteins almost exclusively around the kDNA disc, colocalising with the signal for TAC102 (Figure 2B). Thus, consistent with the localisation of the fusion protein, the biotinylated proteins are mostly in proximity to the kDNA.

**Figure 2.**
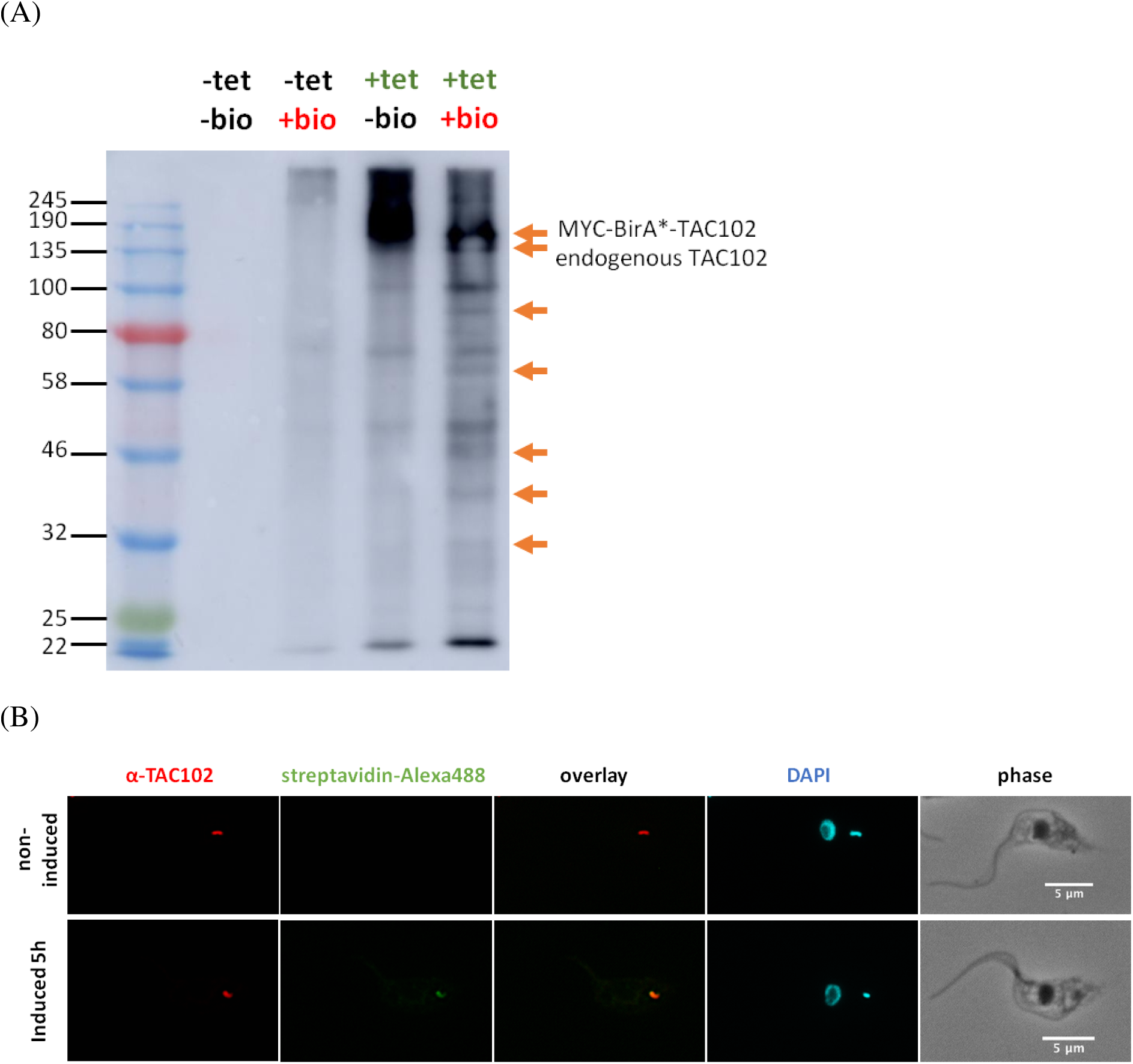
Biotinylation by Myc-BirA*-TAC102 in PCF cells. (*A*) Western blot analysis of biotinylated proteins from the Myc-BirA*-TAC102 cell line (PCF) with (“+ tet”) or without (“-tet”) induction with tetracycline overnight and with (“+ bio”) or without (“-bio”) addition of biotin in the medium. Left lane indicates protein size in kDa. Arrows indicate examples of proteins only detected in the condition “+ tet, + bio”. The two upper arrows indicate the expected size for Myc-BirA*-TAC102 and for the endogenous TAC102 protein. (B) Immunofluorescence microscopy pictures of Myc-BirA*-TAC102 cell line (PCF) with or without induction with tetracycline for five hours. A monoclonal antibody is used to visualise TAC102. Streptavidin-Alexa 488 recognizes the biotinylated proteins. DAPI stains nuclei and kDNA.

TAC102 BioID was performed as described in the supplement (Figure S1A). In brief, biotin was added to procyclic form cells 24 hours prior to the expression of Myc-BirA*-TAC102. After six hours of Myc-BirA*-TAC102 expression the cells were cells were lysed with detergent (0.5% NP-40) in a buffer containing protease inhibitors. After centrifugation the soluble fraction was incubated with streptavidin-conjugated magnetic beads to bind and enrich the biotinylated peptides. After washing the biotinylated peptides were released from the beads by boiling with Laemmli buffer. Protein identification was done by mass spectrometry. The enrichment of biotinylated proteins in cells induced for the expression of Myc-BirA*-TAC102 was compared to non induced cells.

### TAC102 BioID identifies mostly mitochondrial proteins

We analysed the biotinylation pattern induced by Myc-BirA*-TAC102 in four biological replicates. Enrichment in the condition containing tetracycline (“+ tet” condition) compared to the condition without tetracycline (“-tet” condition) was calculated for the detected proteins. A Student t-test was performed to determine the significance of the changes. Based on this analysis TAC102 is the most enriched protein (Figure 3). Overall, 77 proteins were enriched at least three fold (significance p ≤ 0.01) (Table S1). Of these proteins, 47 are predicted to have a mitochondrial localisation. Sixteen of the 47 proteins with predicted mitochondrial localisation are annotated as hypothetical proteins and six of these are in the top ten most enriched proteins. Aside from the hypothetical proteins, we identified 11 translation factors, six RNA binding proteins, five DNA binding proteins, five components of the oxidative phosphorylation machinery, three mitochondrial import factors and two nuclear import/export proteins.

**Figure 3.**
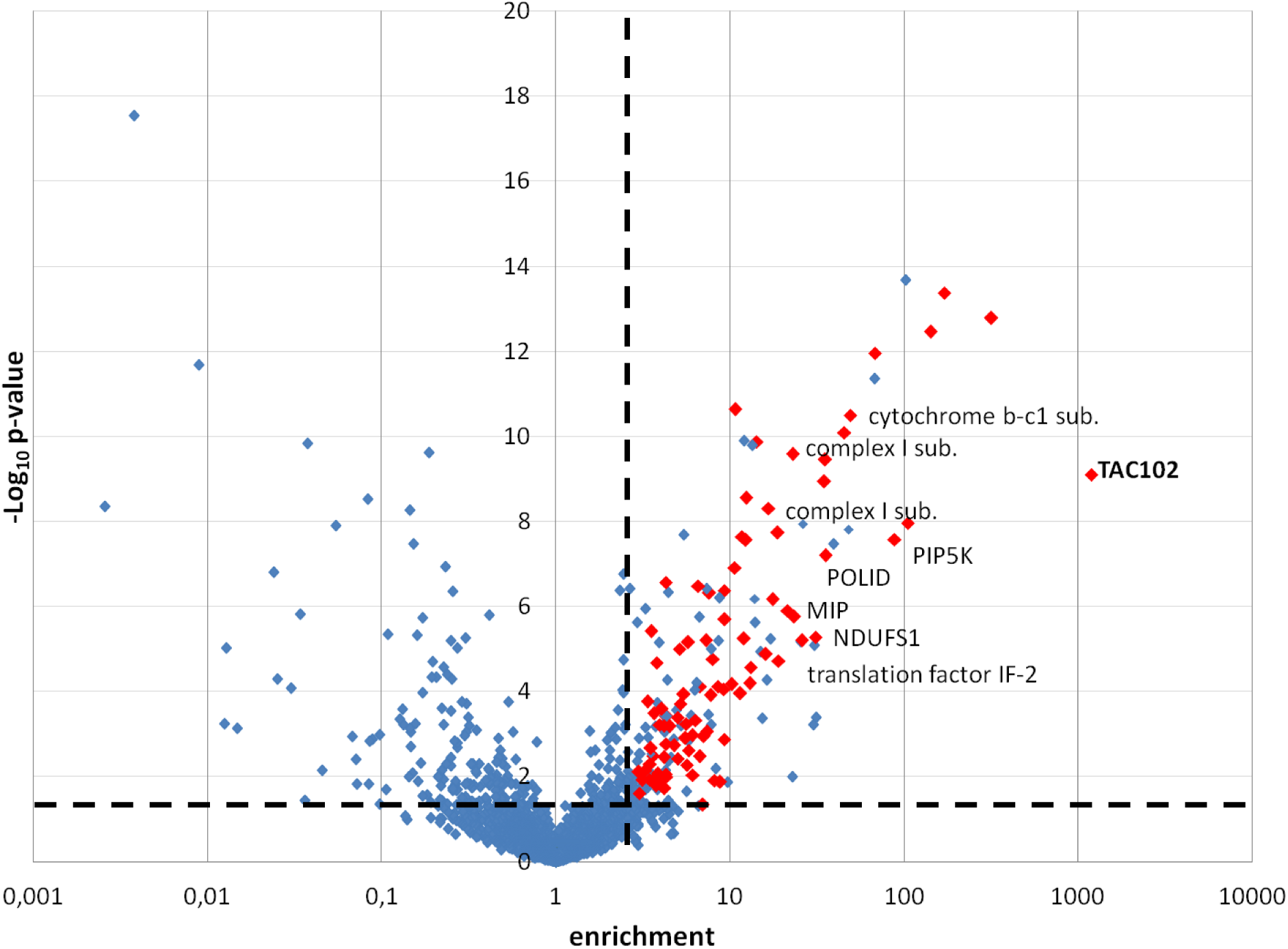
TAC102 BioID enriched proteins. Mitochondrial proteins identified with an enrichment > 3 (p < 0.05) are shown in red. IF: initiation factor; MIP: mitochondrial intermediate peptidase; NDUFS1: NADH-ubiquinone oxidoreductase complex I subunit; PIP5K: phosphatidylinositol-4-phosphate 5-kinase related; POLID: mitochondrial DNA polymerase I D; SSU: small subunit; sub.: subunit; tet: tetracycline.

### TAC102 immunoprecipitation

In order to compare the BioID results with a more conventional approach we used the monoclonal anti-TAC102 antibody coupled to magnetic beads for immunoprecipitation experiments. PCF cells were lysed with digitonin and fractionated into an organellar and cytoplasmic fraction by centrifugation. As shown previously TAC102 is found in the organellar fraction (Trikin et al. 2016). After lysis of this fraction with 1% NP-40, about 50% of TAC102 was detected in the soluble fraction, which subsequently was used for the immunoprecipitation (Figure S2A). The two elution fractions (E1 and E2) were resolved by SDS-PAGE and analysed by western-blot using the monoclonal anti-TAC102 antibody. TAC102 was enriched in the first elution step (E1) (Figure S2B) while the second elution fraction (E2) did not show a detectable amount of TAC102. In the fractions resolved on SDS-PAGE and stained with silver we found the majority of the proteins in the flowthrough (Figure S3). The silver staining of the gel identified a prominent band in the elution (E1 fraction) between 100 kDa and 135 kDa, which likely corresponds to TAC102 itself (Figure S3).

### TAC102 immunoprecipitation identifies TAC components

Immunoprecipitation with and without anti-TAC120 antibody coupled to the beads were done in triplicate. We used label free quantification methods to identify and quantify the peptides (Asara et al. 2008) (Silva et al. 2006; Grossmann et al. 2010). Enrichment was calculated and statistical significance was tested by an empirical Bayes test and the p-value was corrected by the Benjamin and Hochberg false discovery rate method, with a false discovery rate of 0.01. Of the 775 proteins that we detected, 100 proteins were enriched at least three fold in the TAC102 immunoprecipitation (significance p ≤ 0.01) (Figure 4, and Table S2). Of these 100 proteins, 49 are predicted to have a mitochondrial localisation. TAC102 was the most enriched protein followed by p166 another TAC component. Additionally we detected two TAC components of the outer mitochondrial membrane, namely TAC40 and TAC60 among the top ten most enriched proteins. Nine of the 49 proteins with mitochondrial localisation are annotated as hypothetical proteins and four of these are in the top ten most enriched proteins. Aside from the hypothetical proteins and TAC components, we identified 30 translation factors, ten RNA binding proteins, seven DNA binding proteins, six ribosome biogenesis factors, two cristae formation proteins, two flagellum attachment zone proteins, two protein folding factors and two rRNA processing factors.

**Figure 4.**
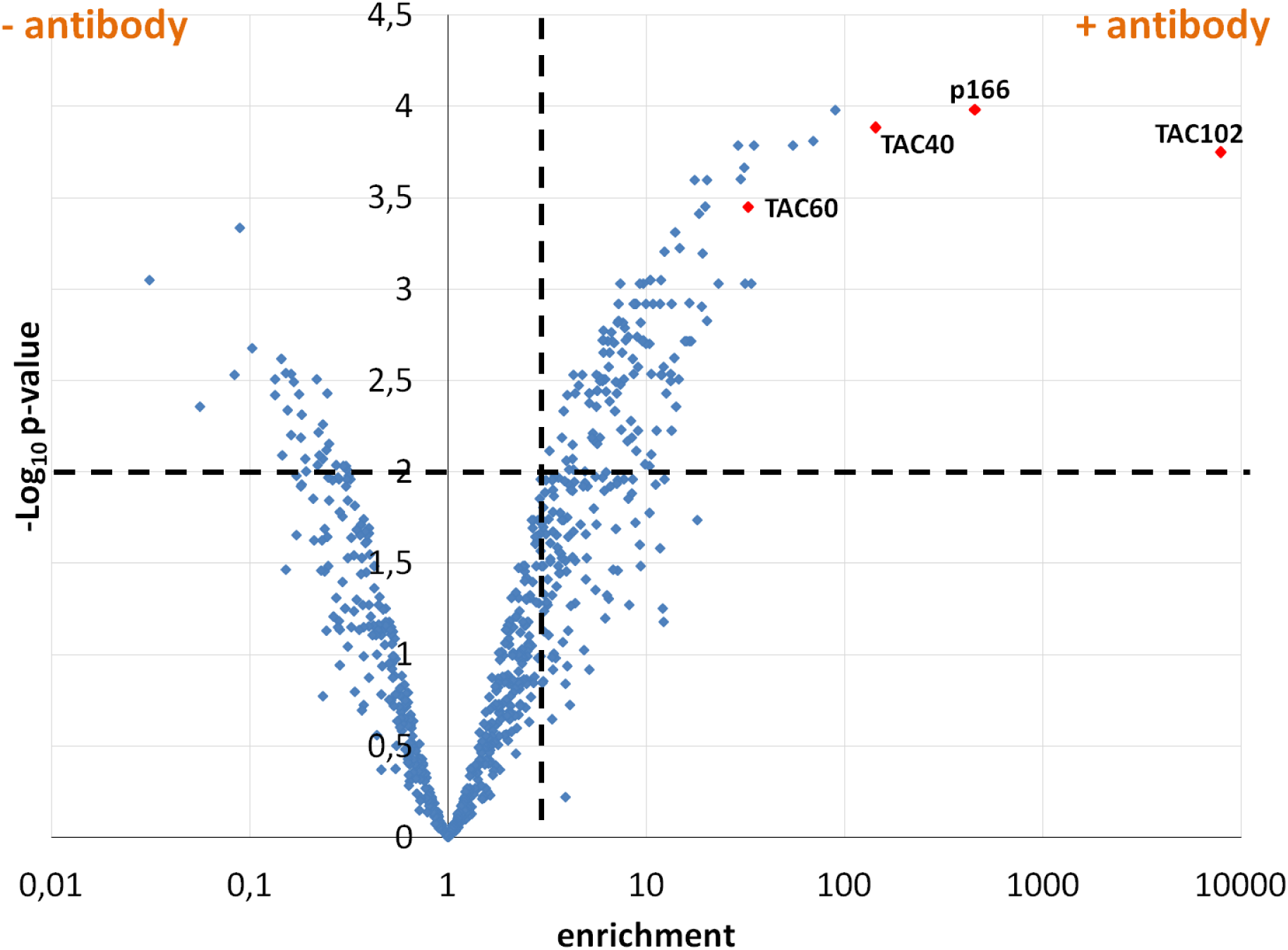
Volcano plot representing the significance of the results from mass spectrometry analysis of TAC102 immunoprecipitation compared to the enrichment. A total of 775 proteins were identified from which 100 proteins are significantly enriched in the TAC102 immunoprecipitation (p ≤ 0.01). The two dotted lines represent the cut-off used (p ≤ 0.01 and enrichment > 3). The protein the most enriched in the condition is TAC102. Three others TAC components were also from the most enriched (p166, TAC40 and TAC60).

When comparing the BioID and TAC102 immunoprecipitation data, we identified nine proteins to be significantly enriched in both approaches (Table 1). Aside from the bait TAC102 these were the mitochondrial protein import receptor ATOM69, two ribosomal proteins and five proteins with unknown function. In order to verify the localisation of the five proteins, we aimed to tag the corresponding genes in PCF trypanosomes *in situ* at the 3’ end with a triple hemagglutinin or a myc tag using a PCR based approach (Oberholzer et al. 2006). Three tagged proteins localised to the kDNA/TAC region (Tb927.10.900, Tb927.8.3160, Tb927.9.6410) one seemed to primarily localise in the cytoplasm (Tb927.7.850) and for one of the candidates we were unable to produce a C-terminally tagged cell line (Tb927.7.5330) (Figure 5).

**Table 1.**
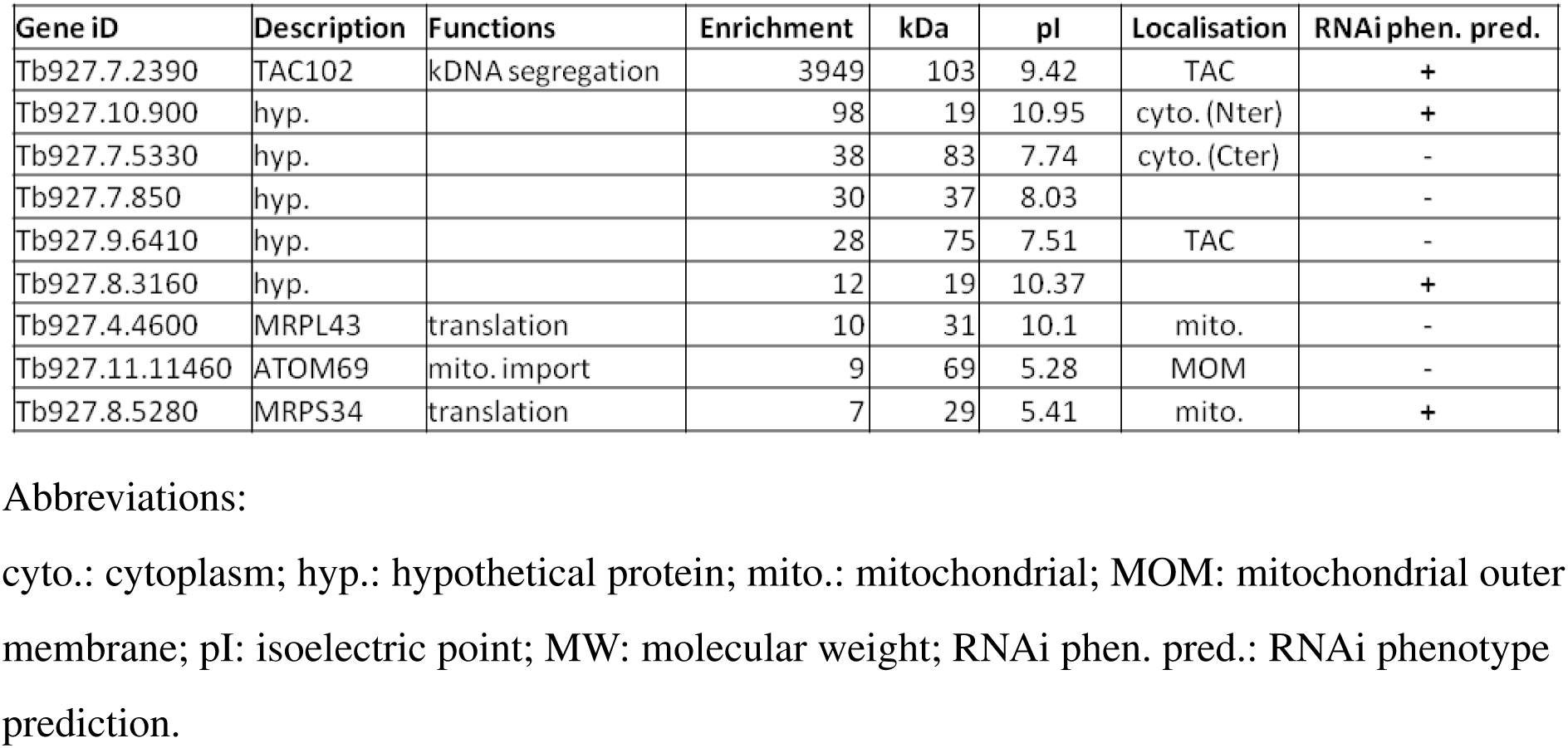
Nine proteins are enriched significantly in both TAC102 BioID and TAC102 immunoprecipitation (enrichment > 3; p ≤ 0.01)

| Gene iD | Description | Functions | Enrichment | kDa | pI | Localisation | RNAi phen. pred. |
| --- | --- | --- | --- | --- | --- | --- | --- |
| Tb927.7.2390 | TAC102 | kDNA segregation | 3949 | 103 | 9.42 | TAC | + |
| Tb927.10.900 | hyp. |  | 98 | 19 | 10.95 | cyto. (Nter) | + |
| Tb927.7.5330 | hyp. |  | 38 | 83 | 7.74 | cyto. (Cter) | - |
| Tb927.7.850 | hyp. |  | 30 | 37 | 8.03 |  | - |
| Tb927.9.6410 | hyp. |  | 28 | 75 | 7.51 | TAC | - |
| Tb927.8.3160 | hyp. |  | 12 | 19 | 10.37 |  | + |
| Tb927.4.4600 | MRPL43 | translation | 10 | 31 | 10.1 | mito. | - |
| Tb927.11.11460 | ATOM69 | mito. import | 9 | 69 | 5.28 | MOM | - |
| Tb927.8.5280 | MRPS34 | translation | 7 | 29 | 5.41 | mito. | + |

Abbreviations:
cyto.: cytoplasm; hyp.: hypothetical protein; mito.: mitochondrial; MOM: mitochondrial outer membrane; pI: isoelectric point; MW: molecular weight; RNAi phen. pred.: RNAi phenotype prediction.

**Figure 5.**
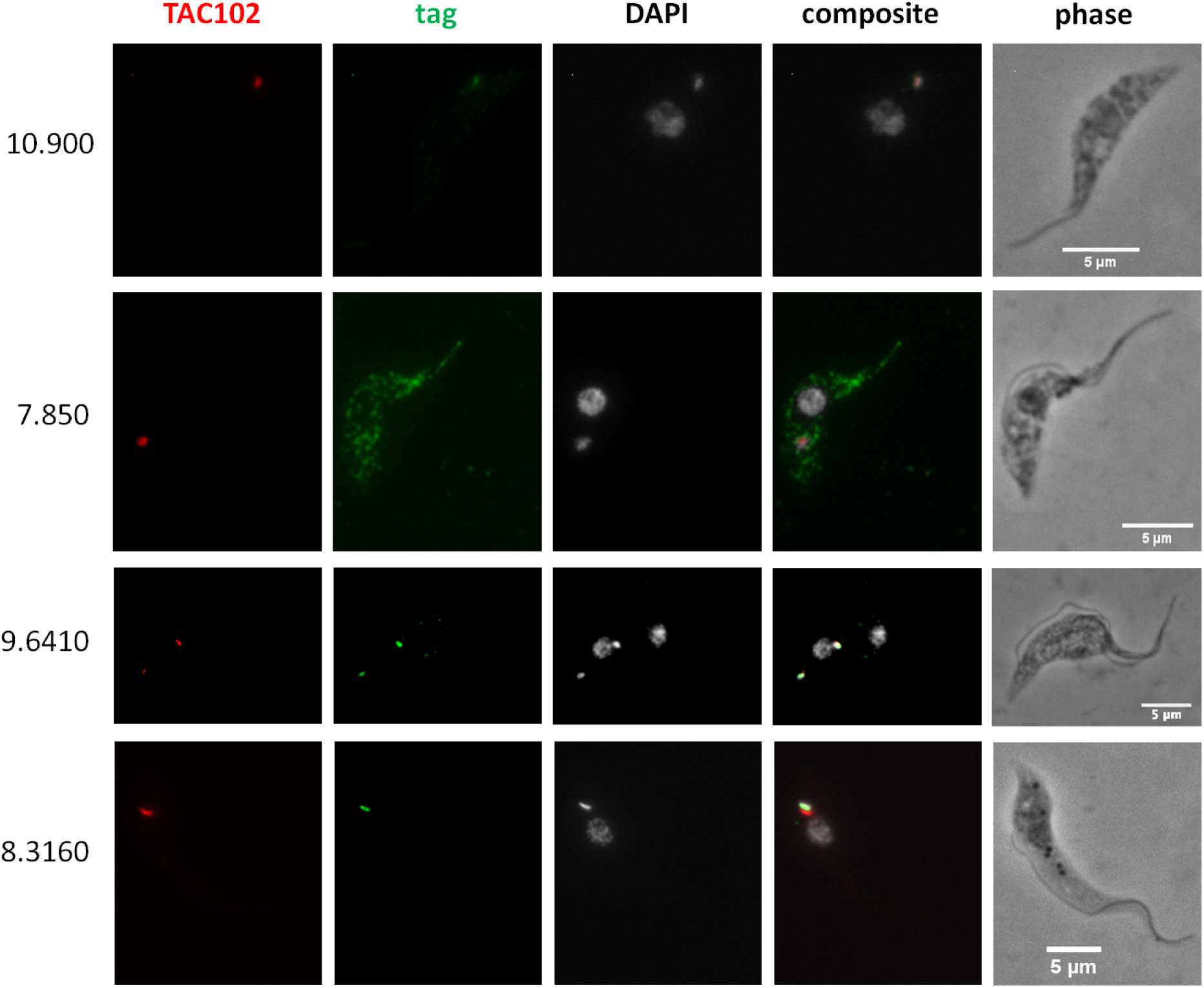
Localisations in procyclic forms (PCF) by immunofluorescence microscopy of the hypothetical proteins enriched significantly in both TAC102 BioID and TAC102 immunoprecipitation. TAC102 is recognised by a monoclonal antibody. The candidate is HA tagged (10.900, 7.850 and 8.3160) or MYC tagged (9.6410) and is recognised by an anti-HA antibody or an anti-MYC antibody. DAPI stains the DNA content (nuclei and kinetoplasts).

### Yeast two-hybrid screen shows interaction between TAC102 and p166

The yeast two-hybrid screen (Hybrigenics) was done essentially as described previously (López-Farfán et al. 2014). The entire open reading frame of TAC102 was fused to the C-terminus of LexA (N-LexA-TAC102) of and expressed in yeast cells. A total of 138 million interactions were screened and 15 clones were isolated and characterised (Table S3). Three high confidence hits were identified expressing a N-terminal part of the TAC component p166 ((Zhao et al. 2008; Hoffmann et al. 2018)). This large, acidic TAC protein contains a single transmembrane domain in the C-terminus and was previously shown by superresolution microscopy to be in close proximity to TAC102 (Zhao et al. 2008; Hoffmann et al. 2018). Two of the three high confidence p166 clones expressed a short region from amino acid 37 to 210 while the third clone expressed a larger fragment ranging from amino acid 71 to 716 of the 1502 amino acid long protein (Figure S4). The only other hits with good confidence were two identical clones of the C-terminal region of the putative nuclear pore component NUP109, that was previously found in a proteomics analysis characterising nuclear proteins (Goos et al. 2017).

## Discussion

In this study we employed three different approaches with the aim to identify interacting proteins of TAC102, which is a key component of the mitochondrial genome segregation machinery. We identified at least three potential novel TAC components and could establish the direct interaction of TAC102 and p166.

One of the approaches applied proximity-dependent biotin identification (BioID), which was previously used by Morriswood and colleagues for the identification of proteins localised in the bilobe region of the flagellum in *T. brucei* (Morriswood et al. 2013). While there are a few studies using BioID in combination with mitochondrially targeted proteins, like the protein interaction study on Clp in human 293T cells (Cole et al. 2015) or the mitochondrial PolG interactome (Liyanage et al. 2017), in this study we show for the first time the potential of BioID with a mitochondrial protein of a protozoan from the group of the Excavata. Although we did not formally test if the Myc-BirA*-TAC102 fusion protein is capable of rescuing a TAC102 knockout, the localisation by immunofluorescence microscopy (see Figure 1A), the biochemical behavior during detergent solubilization (Figure 1B) and the lack of a dominant negative phenotype argue that the protein is likely assembled in the TAC. The detection of multiple signals for TAC102/Myc-BirA*-TAC102 on western blot after the biochemical fractionation is potentially due to the instability of the TAC102 when released from the TAC structure. Alternatively the smaller bands visible on the western blot could also be due to posttranslational modifications. Consistent with the localisation of the fusion construct Myc-BirA*-TAC102 in the mitochondrial organelle, most of the identified interactors (> 60%) are known mitochondrial proteins and half of those are known to localise at the kDNA or the TAC itself (Table S1). While the identification of kDNA associated proteins is not surprising since they are in close proximity to the TAC, we also identified two components of the mitochondrial protein import machinery. One is ATOM69, a receptor of the major import pore, the archaic translocase of the mitochondrial outer membrane (ATOM) in trypanosomes (Mani et al. 2015). In most cases protein import into the mitochondrion requires an N-terminal targeting signal. The Myc-BirA*-TAC102 fusion protein however uses its natural import signal that was previously shown to reside in the C-terminal region of TAC102 (Trikin et al. 2016). Thus unfolding and import from the C-terminus of the fusion construct would allow for biotinylation of the receptor during the import process. The second protein that is part of the import process and seems to interact with Myc-BirA*-TAC102 is the mitochondrial intermediate peptidase MIP, which was previously described to be essential in PCF cells (Peña-Diaz et al. 2018). We asked if depletion of the peptidase would lead to TAC102 precursor accumulation. RNAi targeting the mitochondrially localised MIP in BSF cells led to a very strong growth defect, however no precursor accumulation of TAC102 was observed (Figure S5). Thus, either there is no processing and the interaction of TAC102 and MIP serves a different purpose, or the processed peptide is very short and not readily detectable, or the MIP is a false positive interaction. Compared to the BioID approach, immunoprecipitation using a monoclonal TAC102 antibody provided the advantage of targeting the native protein rather than relying on an artificial fusion protein. However, the immunoprecipitation with TAC102 as bait did not lead to a greater enrichment of mitochondrial proteins, when compared to the BioID approach and in both cases TAC102 was the most highly enriched protein. Interestingly, the TAC102 immunoprecipitation identified the TAC component p166 as the second most enriched protein (after TAC102), while BioID did not detect any currently characterised TAC component except TAC102 itself. The interaction of TAC102 and p166 was confirmed by the yeast two-hybrid screen that demonstrated the TAC102 interaction domains are in the N-terminal region of p166 (Figure S4, Table S3). Thus the C-terminus of p166 with its single predicted transmembrane domain is likely to reside at the inner mitochondrial membrane, while the N-terminus connects to the kDNA proximal TAC102. So, if p166 is a direct interactor of TAC102, why did the immunoprecipitation, but not the BioID approach identify this interaction? One explanation could be the orientation of the BirA*-TAC102 fusion protein in the TAC. If TAC102 is interacting with p166 via its C-terminus, then the BirA* moiety of the fusion protein might be facing the kDNA and be too far away from p166 for proximity labeling. This would also explain why the BioID approach identified seven proteins close or in the kDNA network. Alternatively, the BirA*-TAC102 fusion protein, despite its apparent correct localisation, might not be incorporated properly into the TAC and thus not be in proximity to p166. Aside from p166, the TAC102 immunoprecipitation also enriched two TAC components of the outer mitochondrial membrane (TAC40 and TAC60, Figure 6), which are in close proximity to p166 in the TAC (Hoffmann et al. 2018). Both approaches (BioID and IP) did enrich a known component of the kDNA replication machinery (Figure 6). TAC102-BioID identified the mitochondrial PolID, which dynamically localises at the antipodal sites during minicircle replication (Concepcion-Acevedo, Luo, and Klingbeil 2012). TAC102 immunoprecipitation identified Pol beta PAK a polymerase localised throughout the kDNA disc, which is believed to be involved in minicircle gap closure prior to segregation of the network (Saxowsky, Choudhary, and Klingbeil 2003). Since replication of the minicircles occurs in the kinetoflagellar zone which is also the home of the TAC, it is not surprising to find at least some of the replication components interacting with the segregation machinery (Figure 6). In conclusion we have for the first time used BioID to identify intractors of a mitochondrial protein in trypanosomes and compared the approach to immunoprecipiation and yeast two-hybrid. We were able to identify three proteins that by their localization could potentially represent novel TAC components and established the direct interaction of TAC102 with the N-terminal region of p166.

**Figure 6.**
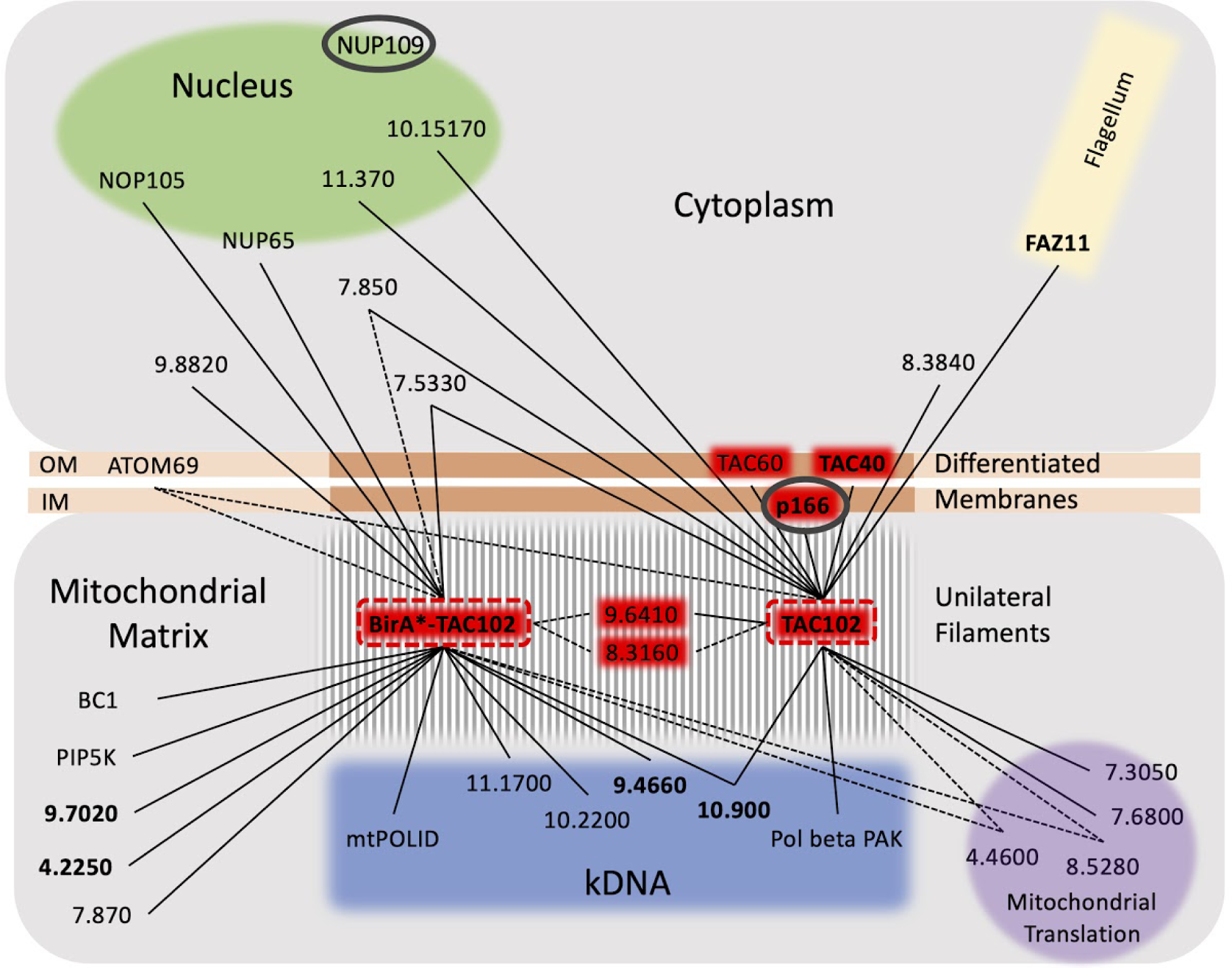
Depiction of the TAC102 interactions as identified from BioID, immunoprecipitation and yeast two hybrid screen. The common interactors (BioID/CoIP) are connected with a dotted line (-----). The 15 most enriched proteins of the TAC102 IP or BirA*-TAC102 (BioID) are connected with a solid line (___). The five most enriched proteins are in **bold**. The two proteins identified by the yeast two hybrid screen are circled in grey (∘). BC1: cytochrome b-c1 subunit; IM: inner mitochondrial membrane; NOP: nucleolar protein; NUP: nucleoporin; OM: outer mitochondrial membrane; PIP5K: phosphatidylinositol-4-phosphate 5-kinase related; POLID: DNA polymerase I D; Pol beta PAK: Polymerase beta with PAK domain.

## Material and methods

### Antibodies and reagents

The rabbit polyclonal anti-Myc antibody and the Biotin was purchased from Sigma-Aldrich. The mouse anti-EF1α antibody was purchased from Santa Cruz Biotechnology. The mouse monoclonal anti-TAC102 antibody and the rabbit anti-ATOM antibody have been described previously (Trikin et al. 2016; Pusnik et al. 2011). Streptavidin-conjugated magnetic beads were purchased from NEB. HRP-conjugated streptavidin and AlexaFluor488-conjugated streptavidin were purchased from Thermo Fisher Scientific.

### Trypanosomes, culture and generation

We cultured procyclic form (PCF) Lister 427 29-13 and bloodstream form (BSF) Lister 427 SM, *Trypanosoma brucei brucei* as described previously (Amodeo, Jakob, and Ochsenreiter 2018). In brief, PCF/BSF cells were grown at 27°C/37°C without and with 5% (v/v) CO_2_, in SDM79/HMI9 medium supplemented with 10% (v/v) of fetal calf serum (Sigma-Aldrich). Myc-BirA*-TAC102 expression was induced by tetracycline (1 µg/ml).

### Wide field fluorescence microscopy

5×10^5^ cells were allowed to settle on a slide for 40 min. The cells are fixed for four minutes with 4% (m/v) paraformaldehyde (PFA 4%) in PBS. Cells are then permeabilized with 0.2% (v/v) Triton X-100 for five minutes. After each treatment, the slide is washed with PBS. The slide was then blocked in PBS-BSA 4% (m/v) for 30 min in a humid atmosphere. The primary antibodies and secondary antibodies are added to the slide and incubated for one hour each at room temperature. The primary and secondary antibodies are diluted in PBS-BSA 4% (anti-Myc rabbit 1/1000, anti-TAC102 mouse 1/5000). Post staining cells are mounted in ProLong^®^ Gold antifade reagent (Life technologies) containing 4′,6-diamidino-2-phenylindole (DAPI).

The slides are observed with a 100x oil immersion phase contrast objective on the Leica DM 5500 fluorescence microscope.

### BioID

The primers used for the amplification of TAC102 open reading frame are 2390 BamHI fwd: 5’-CGGGATCCATGTATCGGCCTCGTGGCGG-3’ 2390 SalI rev: 5’-CGGGTCGACTTACTTTATAAGCTGCCGAA-3’

The sequence coding for TAC102 is cloned in between the restriction sites of the enzymes *Xho*I and *BamH*I into the pLew100_Myc_BirA vector (Morriswood et al. 2013). The protocol used is an adaptation of the one described in (Morriswood et al. 2013). 2.5 ml of biotin stock solution (1 mM in sterile MilliQ water) is added to 47.5 ml of PCF cells and incubated for 24 h, in order to give biotin time to enter the cells. The expression of Myc-BirA*-TAC102 is subsequently induced for six hours with tetracyclin. We used 5×10^8^ cells per experiment. They were collected by centrifugation (1800 g, 5 min, 4°C), washed three times with phosphate-buffered saline (PBS: 137 mM NaCl, 2.7 mM KCl, 10 mM Na_2_HPO_4_, 2 mM KH_2_PO_4_) and finally resuspended in PEME buffer (2 mM EGTA, 1 mM MgSO_4_, 0.1 mM EDTA, 0.1 M PIPES, pH 6.9) containing 0.5% (v/v) NP-40 and protease inhibitors (cOmplete ULTRA Tablets, Mini, EDTA-free, Roche). The tube is left for 15 min at room temperature with gentle mixing: this is the E1 fraction (see Figure S1A). The tube is then centrifuged (3400xg, 2 min, RT) and the supernatant is put in a new tube: this is the S1 fraction. The pellet is then resuspended in lysis buffer (0.4% (m/v) SDS, 500 mM NaCl, 5 mM EDTA, 1 mM DTT, 50 mM Tris-HCl pH 7.4). The tube is left for 30 min at room temperature with a gentle mixing: this is the P1 fraction. After centrifugation (16 000 g, 10 min, RT), the supernatant is put in a new tube: this is the S2 fraction. 250 µL of streptavidin conjugated magnetic beads (NEB) are added to S1 and S2. The tubes are then incubated for four hours at 4°C before separation of the beads from the liquid with a magnet: these are fractions F1 and F2. The beads are then washed twice with PBS, centrifuged (6000 g, 2 min, RT) and finally resuspended in Laemmli buffer (5X stock solution: 2% (m/v) SDS, 60 mM Tris-HCl pH 6.8, 24% (v/v) glycerol, 5% (v/v) β-mercaptoethanol, bromophenol blue). Fractions from each sample were separated by SDS-PAGE, transferred to polyvinylidene difluoride (PVDF) membrane and blotted with HRP-conjugated streptavidin.

### Immunoprecipitation

For the immunoprecipitation of TAC102, anti-TAC102 antibody is crosslinked to protein A / protein G beads from the Pierce™ Crosslink Magnetic IP kit (Thermo Scientific™). A mitochondrial enriched fraction is prepared by digitonin fractionation with 100 ml of procyclic form parasites (cell line: 29.13) grown in the exponential phase. The mitochondrial enriched fraction is then resuspended in ice-cold IP lysis/wash buffer (25 mM Tris, 150 mM NaCl, 1 mM EDTA, 1% (v/v) NP-40, 5% (v/v) glycerol) containing protease inhibitors. The sample is incubated on ice for five minutes, with vortexing every minute. After centrifugation (13 000 g, 10 min, 4°C), the supernatant is kept and put on beads containing anti-TAC102 monoclonal antibodies or empty beads (for negative controls), for 1:35 h at room temperature with rotation. The mixture lysate/beads is vortexed every 15 min during this incubation. The beads are collected on a magnetic stand for one minute and the flow-through (FT) is kept on ice in a new tube. The beads are washed twice with ice-cold IP lysis/wash buffer containing protease inhibitors and once with ice-cold water containing protease inhibitors. The proteins bound to the beads are then eluted twice using 300 µL of 0.1 M glycine pH 2.4 containing protease inhibitors with a five minutes incubation on a rotating platform. The elutions are saved on ice in new tubes and the pH is neutralised using 30 µL of neutralisation buffer from the kit. In order to get a reasonable amount of proteins to load on a gel for further mass spectrometry analysis, the elutions from immunoprecipitations are acetone precipitated. To do so, four times the sample volume of cold acetone (−20°C) is added into the elutions. The tube is vortexed and incubated for at least one hour at −20°C. After centrifugation (15 000 g, 10 min, 4°C), the supernatant is removed and the rest of the acetone is allowed to evaporate for 30 min at room temperature. The protein pellet is then resuspended in 20 µL of 1x Laemmli buffer for running on SDS-PAGE.

### Yeast two-hybrid (Y2H)

Bait cloning and Y2H screening were performed by Hybrigenics Services SAS, France (http://hybrigenics.com/services). The coding sequence for full-length TAC102 (Tb927.7.2390) was cloned into a plasmid pB27 as a C-terminal fusion to LexA (N-Lex-TAC102). The construct was used as a bait to screen at saturation a highly complex Treu927 genomic fragment library of *T. brucei* constructed into pP6. pB27 and pP6 derive from the original pBTM116 and pGADGH plasmids, respectively. 138 million clones were screened using a mating approach with Y187 (matα) and L40^Gal4 (mata) yeast strains as previously described (Fromont-Racine, Rain, and Legrain 1997). 15 His+ colonies were selected on a medium lacking tryptophan, leucine and histidine. The prey fragments of the positive clones were amplified by PCR and sequenced. The resulting sequences were used to identify the corresponding interacting proteins in *Trypanosoma brucei* Treu927 genome sequence using a fully automated procedure in the GenBank database (NCBI). A confidence score (PBS, for Predicted biological score) was attributed to each interaction as previously described (Formstecher et al. 2005). This global score represents the probability of an interaction being nonspecific. The PBS scores have been used to select out of the 15 clones those with high confidence interaction (Score B).

Biochemical methods including digitonin fractionation, coomassie and silver stained gels were done as described previously (Amodeo, Jakob, and Ochsenreiter 2018). For mass spectrometry analysis the samples were briefly separated on precast 10% polyacrylaminde gels under denaturing conditions. Pieces of the gel were then treated essentially as described previously (Gunasekera et al. 2012).

## End section

### Data, code and materials

The datasets supporting this article have been uploaded as part of the supplementary material.

### Authors’ contributions

HB carried out the biochemical lab work, participated in the molecular lab work, cell culture, immunofluorescence, design of the study, data analysis and drafted the manuscript; LP participated in the molecular lab work, cell culture and immunofluorescence; TO coordinated the study, participated in data analysis, design of the study and revised the manuscript. All authors gave final approval for publication and agree to be held accountable for the work performed therein.

### Funding

This work was supported by the Swiss National Science Foundation (17423) and the canton of Bern.

## Acknowledgments

We thank Roman Trikin and Ana Kalichava for technical assistance, Simona Amodeo and Irina Bregy for reading the manuscript. We thank Andre Schneider and Volker Heussler for antibodies. We also thank the Proteomic and Mass Spectrometry Core Facilities (PMSCF) and the Microscopy Imaging Centre (MIC) from the University of Bern, Switzerland.

