## Supplemental material for "Discovery of new mitochondrial DNA segregation machinery components in *Trypanosoma brucei*: a comparison of three approaches"

**Supplementary information**

(A)

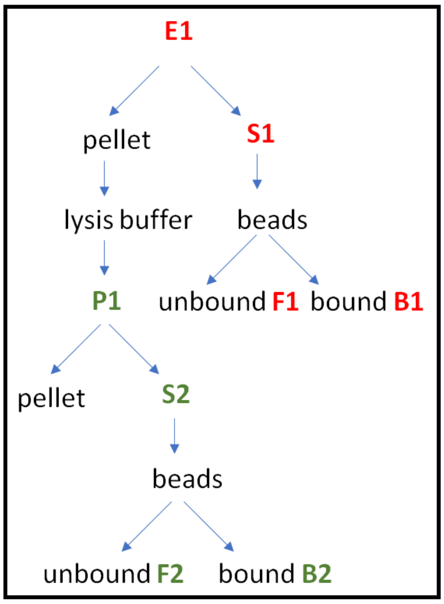


(B)

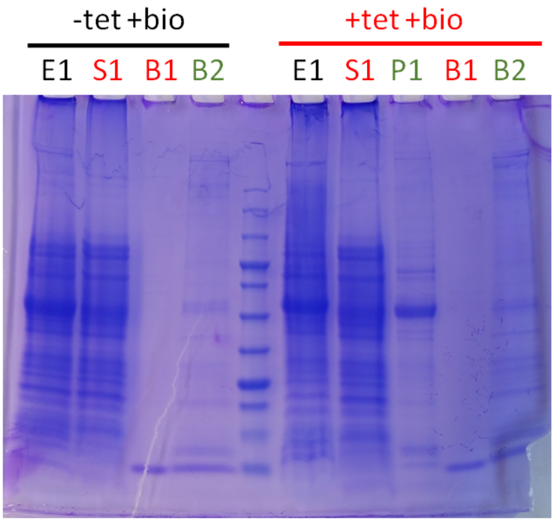


(C)

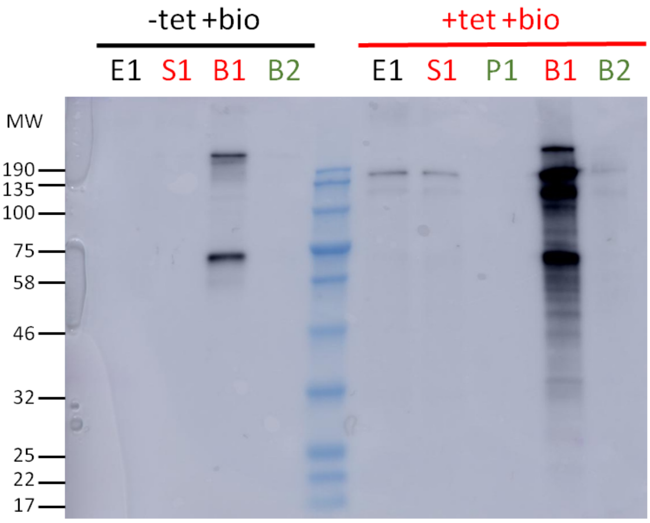


**Figure S1.** Purification of TAC102 binding partners and near neighbours using BioID. (*A*) Schematic of purification protocol. (*B*) Coomassie staining of the SDS-PAGE gel from BioID fractions. (*C*) Western blot analysis of the BioID fractions. Total cells extracts (E1), cleared supernatants (S1), first pellet (P1) and streptavidin bead bound (B1, B2) fractions were loaded on the gel. For E1, S1 and P1, the equivalence of 7.2x$10^6$ cells is loaded. For B1 and B2, the equivalence of 137x$10^6$ cells is loaded. MW: molecular weight.

(A)

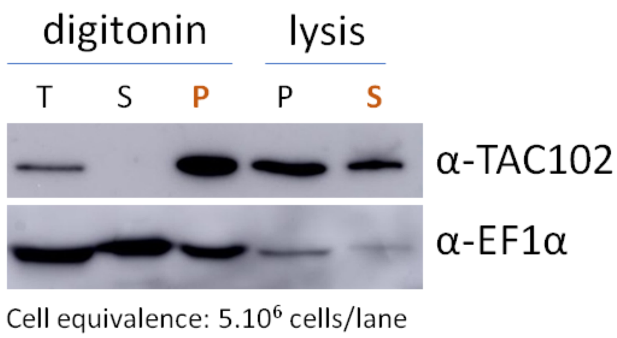


(B)

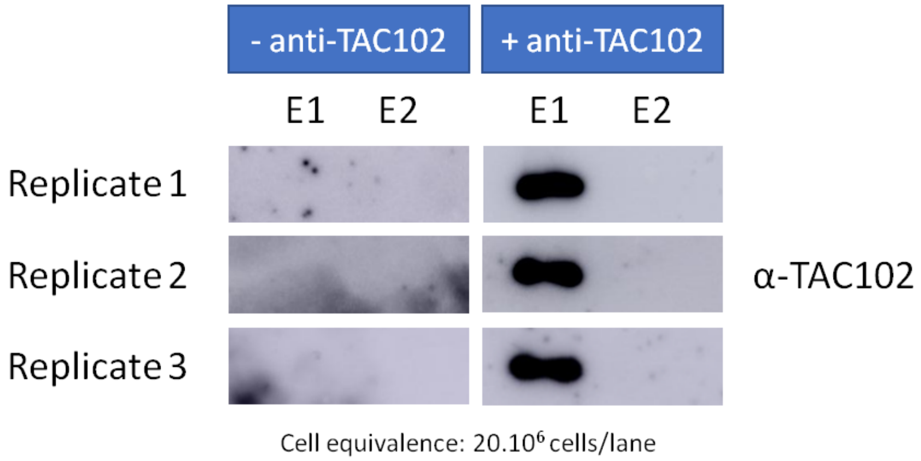


**Figure S2.** TAC102 enrichment and immunoprecipitation. (*A*) Western blot analysis of the digitonin fractionation and subsequent lysis of the pellet fraction with 1% (v/v) NP-40 of whole cell protein from procyclic cells. T: total; S: supernatant; P: pellet. In orange: the pellet from the digitonin fractionation is used for the subsequent NP-40 lysis. The elongation factor EF1α serves as a control (*B*) Western blot analysis of the elutions (E1 and E2) of TAC102 immunoprecipitation replicates.

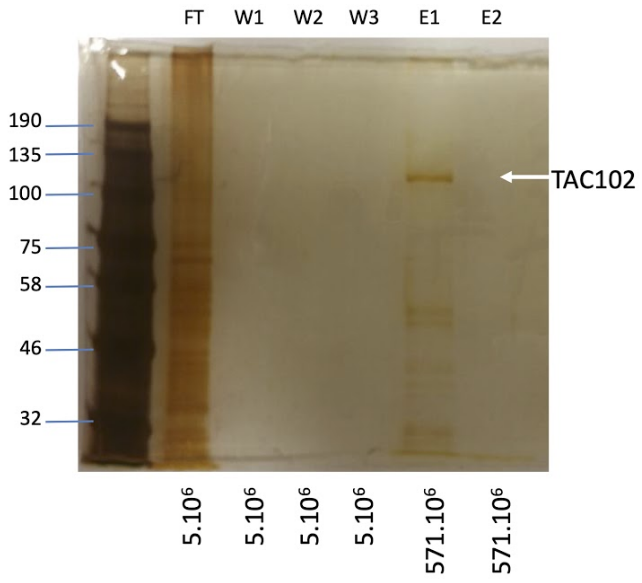


**Figure S3.** Silver stained SDS PAGE of TAC102 immunoprecipitation fractions. T: total S: supernatant; P: pellet; TAC102: TAC component (mitochondrion); EF1α: Elongation Factor 1 α: translation elongation factor (cytosol).

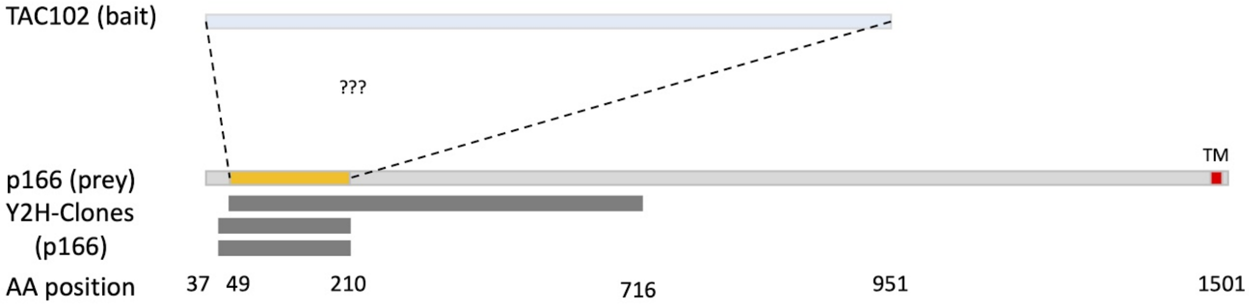


**Figure S4.** TAC102 yeast two-hybrid screen high confidence interactions. Depicted are the three high confidence interaction clones (Y2H-Clones, dark grey) that express a N-terminal region of p166. p166 (light grey) is shown as reference including the C-terminal predicted transmembrane (TM) domain. The region sufficient for TAC102 interaction is shown in yellow. (???) The question marks symbolize that the domains within TAC102 responsible for binding to p166 are unknown.

(A)

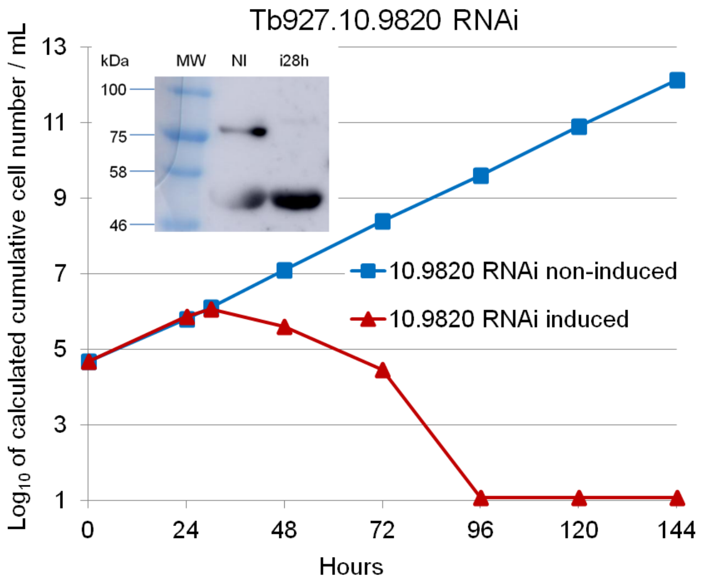


(B)

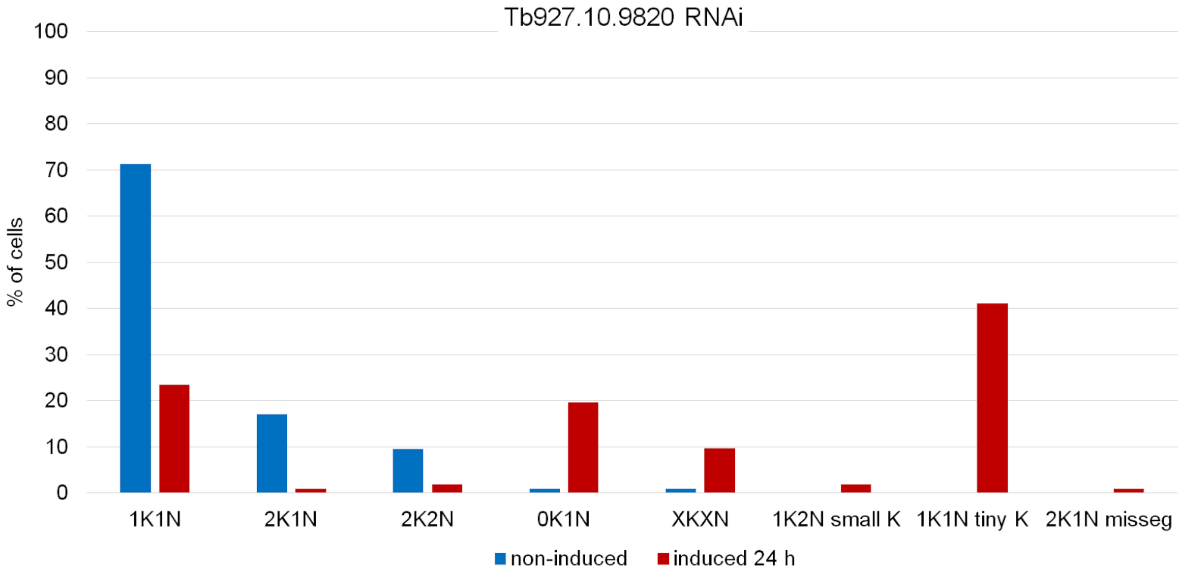


(C)

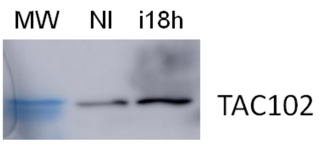


3.5% gel

(D)

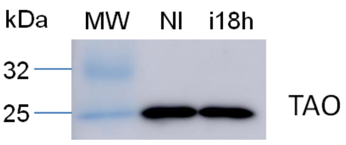


12% gel

(E)

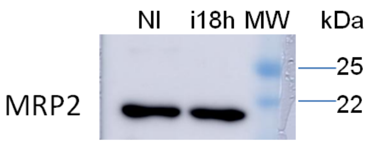


12% gel

**Figure S5.** Characterisation of MIP RNAi cell line (BSF) (gene: Tb927.10.9820). (*A*) Growth curves of MIP RNAi cell line with induction (red) and without induction (blue) with tetracycline. MIP has been C-terminally Myc-tagged in the RNAi cell line. The fusion protein is expected at 80 kDa. The RNAi efficiency was assessed by western blot using an anti-Myc antibody (rabbit). The upper band corresponds to MIP-Myc. The lower band is an unspecific band detected by the antibody. (*B*) Cell cycle status of MIP RNAi cells (BSF) with (red) and without (blue) induction of RNAi for 24 h. Around 100 cells are observed per condition (non-induced and induced). (*C*) Western blot of MIP RNAi cell line probed for TAC102. A 3.5% gel was used. (*D*) Western blot of MIP RNAi cell line probed for TAO. A 12% gel was used. (*E*) Western blot of MIP RNAi cell line probed for MRP2. A 12% gel was used. kDa: kilodalton; MW: molecular weight; NI: RNAi non-induced; i28h: RNAi induced during 28 h.

**Table S1.** Enriched Proteins from Myc-BirA*-TAC102 BioID (enrichment > 2 ; p ≤ 0.01)

| Gene ID | Description | Functions | Enrichment | kDa | Iso. p. | Localisation |
|---|---|---|---|---|---|---|
| Tb427.07.2390 | TAC102 | kDNA segregation | 1192 | 103 | 9.34 | TAC |
| Tb927.9.4660 | hyp. | | 316 | 56 | 5.8 | kDNA |
| Tb427tmp.160.5110 | hyp. (Tb927.9.7020) | | 170 | 25 | 4.83 | mito. |
| Tb427.04.2250 | hyp. | | 142 | 39 | 6 | mito. |
| Tb927.10.900 | hyp. | | 105 | 19 | 10.95 | mito. |
| Tb927.10.3810 | NUP65 | nuclear import / export | 102 | 65 | 8.83 | nucl. |
| Tb927.1.740 | PIP5K | lipid kinase act. | 88 | 49 | 5.51 | mito. |
| Tb927.11.1700 | hyp. | | 68 | 62 | 3.99 | kDNA |
| Tb927.7.650 | NOP105 | nucleolar protein | 67 | 106 | 4.5 | nucleol. |
| Tb427tmp.01.7900 | hyp. (Tb927.11.16220: c. b-c1 s.u., put.) | ox. phos. | 49 | 22 | 9.92 | mito. |
| Tb427.07.870 | hyp. | | 48 | 41 | 7.26 | mito. |
| Tb927.7.5330 | hyp. | | 45 | 83 | 7.74 | cyto. |
| Tb927.9.8820 | hyp. | | 39 | 120 | 9.03 | cyto. |
| Tb427tmp.02.0770 | POLID, put. (Tb927.11.3260) | DNA-directed DNA pol. act. | 36 | 183 | 8.1 | mito. |
| Tb927.10.2200 | hyp. | telomeric DNA bind. | 35 | 38 | 8.98 | kDNA |
| Tb927.8.1500 | hyp. | mRNA bind. | 35 | 63 | 9.03 | |
| Tb927.10.12540 | NDUFS1, put. | glutathione-disulfide reductase act. | 31 | 34 | 5.08 | mito. |
| Tb427.10.3320 | hyp. | | 30 | 61 | 8.02 | cyto. |
| Tb927.7.850 | hyp. | | 26 | 37 | 8.03 | |
| Tb927.2.5050 | prot. phosphatase 2C, put. | phosphatase act. | 26 | 35 | 6.39 | cyto. |
| Tb427.07.3280 | translation IF-2, put. | translation | 25 | 80 | 6.6 | mito. |
| Tb427.10.9820 | MIP | mito. import | 23 | 77 | 6.44 | mito. |
| Tb927.7.7330 | NDUF, put. | ox. phos. | 23 | 47 | 6.27 | mito. |
| Tb927.9.6410 | hyp. | | 21 | 75 | 7.51 | TAC |
| Tb927.7.3810 | Cold-shock' prot., put. | mRNA bind. | 19 | 42 | 5.79 | kDNA |
| Tb927.11.13910 | NDUF, put. | ox. phos. | 19 | 29 | 6.35 | mito. |
| Tb927.9.11880 | mt SSU r. prot., put. | translation | 18 | 51 | 8.83 | mito. |
| Tb927.8.1490 | Prot. (DUF1674), put. | | 17 | 11 | 6.24 | mito. |
| Tb927.8.3160 | hyp. | | 17 | 19 | 10.37 | TAC |
| Tb927.11.9890 | SRP receptor α su., put. | mito. import | 16 | 64 | 5.54 | ER |
| Tb927.10.2800 | MRP s6 | translation | 16 | 19 | 9.31 | mito. |
| Tb927.8.3230 | hyp. | | 15 | 65 | 9.16 | mito. |
| Tb927.1.1530 | Ser/Thr kinase, put. | protein kinase act. | 14 | 191 | 6.11 | cyto. |
| Tb927.7.4620 | hyp. | | 14 | 33 | 10.01 | |
| Tb427.06.1890 | hyp. | | 14 | 69 | 7.64 | cyto. |
| Tb927.7.750 | hyp. | | 13 | 93 | 9.08 | BB |
| Tb427.07.7010 | hyp. | | 13 | 18 | 11.17 | mito. |
| Tb927.11.11380 | SIM prot. | DNA bind. | 13 | 62 | 6.36 | kDNA |
| Tb927.11.6660 | Tex-like prot., put. | DNA bind. | 12 | 104 | 9.85 | kDNA + nucleol. |
| Tb927.3.3540 | NUP53b | nuclear import / export | 12 | 53 | 6.96 | nucl. |
| Tb927.11.11790 | R3H cont. prot., put. | mRNA bind. | 12 | 61 | 5.34 | nucl. |
| Tb927.3.970 | mt SSU r. prot., put. | translation | 12 | 40 | 6.09 | kDNA |
| Tb927.10.11870 | GRBC5 | RNA bind. | 12 | 35 | 7.21 | kDNA |
| Tb927.11.16870 | NDUFA2 | ox. phos. | 11 | 19 | 4.65 | kDNA |
| Tb927.8.7260 | KAP, put. | phosphate ion bind.;struct. mol. act. | 11 | 114 | 10.33 | mito. |
| Tb927.3.4030 | hyp. | mRNA bind. | 11 | 135 | 6.46 | kDNA |
| Tb927.10.15790 | CHCH cont. prot., put. | | 10 | 17 | 7.29 | cyto. |
| Tb927.2.5660 | ADKA, put. | adenylate kinase act. | 9 | 29 | 5.7 | flagel. |
| Tb927.11.11460 | ATOM69 | mito. import | 9 | 69 | 5.28 | mito. |
| Tb927.8.5280 | MRPS34 | translation | 9 | 29 | 5.41 | mito. |
| Tb927.6.4510 | hyp. | | 9 | 37 | 6.92 | mito. |
| Tb927.11.16050 | CS /TPR repeat, put. | protein complex assembly | 9 | 69 | 5.56 | cyto. |
| Tb927.10.12410 | hyp. | | 9 | 25 | 5.1 | |
| Tb927.10.5820 | pr. RanGDP b. protein | import | 8 | 109 | 6.39 | kDNA |
| Tb427.08.3310 | acetyltransferase, put. | histone modification | 8 | 76 | 6.75 | nucl. |
| Tb927.1.3950 | ALAT | aminotransferase act. | 8 | 56 | 7.04 | kDNA |
| Tb927.6.1410 | NDUF, put. | ox. phos. | 8 | 27 | 9.29 | kDNA + mito. |
| Tb927.2.2210 | hyp. | | 7 | 113 | 5.92 | mito. |
| Tb927.4.4600 | MRPL43 | translation | 7 | 31 | 10.1 | mito. |
| Tb927.3.2670 | hyp. | | 7 | 29 | 6.06 | kDNA |
| Tb927.1.4480 | DNA-b. prot., put. | DNA bind. | 7 | 158 | 6.17 | nucl. |
| Tb427tmp.01.6620 | hyp. (Tb927.11.14980: MRPL38) | translation | 7 | 58 | 6.99 | kDNA + mito. |
| Tb927.10.8210 | KREPA2 | RNA editing | 6 | 63 | 7.09 | kDNA |
| Tb427.10.3330 | hyp. | | 6 | 40 | 5.36 | ER |
| Tb927.11.2530 | mt SSU r. prot., put. | translation | 6 | 85 | 9.24 | kDNA + mito. |
| Tb927.10.11260 | mt SSU r. prot., put. | translation | 5 | 22 | 9.82 | kDNA + mito. |
| Tb927.10.12370 | γ-glutamylcysteine synth. | glutathion synthase | 5 | 77 | 6.47 | nucl. |
| Tb927.9.11120 | CPC prot. INCENP N ter., put. | mitosis | 5 | 73 | 9.69 | mito. + axoneme |
| Tb927.10.11080 | PFR prot. | struct. flagellar protein | 4 | 134 | 3.98 | flagel. + BB |
| Tb427tmp.01.8100 | enolase, put. (Tb927.11.16410) | phosphopyruvate hydratase act. | 4 | 56 | 5.92 | cyto. + axoneme |
| Tb427.10.4220 | hyp. | telomeric DNA bind. | 4 | 34 | 7.15 | |
| Tb927.2.5270 | dynein heavy chain, put. | transport | 4 | 486 | 5.27 | axoneme |
| Tb927.11.5780 | mt DNA-directed RNA pol. | mt DNA-directed RNA pol. act. | 4 | 144 | 7.09 | cyto. |
| Tb927.5.2070 | MRPL su., put. | translation | 4 | 69 | 6.26 | mito. |
| Tb927.5.4080 | hyp. | | 3 | 58 | 6.5 | cyto. |
| Tb927.11.13280 | mt RBP2 | RNA bind.;mRNA bind. | 3 | 25 | 9.74 | kDNA |
| Tb927.8.3170 | kDNA r. prot. 9 | translation | 3 | 90 | 8.22 | kDNA |
| Tb927.8.4240 | hyp. | | 2 | 184 | 8.27 | kDNA |
| Tb927.7.2390 | TAC102 | kDNA segregation | 2 | 103 | 9.42 | TAC |

Abbreviations:

ADKA: adenylate kinase; act.: activity; ALAT: alanine aminotransferase; BB: basal body; bind.: binding; c. b-c1 su.: cytochrome b-c1 subunit; cont.: containing; CPC: chromosomal passenger complex; DUF: domain of unknown function; ER: endoplasmic reticulum; flagel.: flagellar; GRBC: guide-RNA binding complex; hyp.: hypothetical protein; IF: initiation factor; iso. p.: isoelectric point; KAP: kDNA-associated protein MIP: mitochondrial intermediate peptidase; mito.: mitochondrial; mol.: molecule; MRP: mitochondrial ribosomal protein; mt. SSU r.: mitochondrial small subunit ribosomal; NDUF: NADH-ubiquinone oxidoreductase subunit; NOP: nucleolar protein; nucl.: nucleus; nucleol.: nucleolus; NUP: nucleoporine; PIP5K: phosphatidylinositol-4-phosphate 5-kinase related; POLID: mitochondrial DNA polymerase I D; pr. RanGDP b.: predicted RanGDP binding; prot.: protein; put: putative; SIM: SUMO-interacting motif; SRP: signal recognition particle; struct.: structural; ter.: terminal

**Table S2.** Proteins enriched in TAC102 immunoprecipitation (enrichment > 2 ; p ≤ 0.01)

| Gene ID | Description | Functions | Enrichment | kDa | Iso. p. | Localisation |
|---|---|---|---|---|---|---|
| Tb927.7.2390 | TAC102 | kDNA segregation | 7895 | 103 | 9.42 | TAC |
| Tb927.11.3290 | p166 | kDNA segregation | 457 | 166 | 5.13 | TAC |
| Tb927.4.1610 | TAC40 | kDNA segregation | 143 | 40 | 7.1 | TAC |
| Tb927.10.900 | hyp. | | 90 | 19 | 10.95 | mito. |
| Tb927.4.5340 | FAZ11 | protein complex bind. | 70 | 95 | 7.13 | FAZ |
| Tb927.9.6410 | hyp. | | 35 | 75 | 7.51 | TAC |
| Tb927.7.850 | hyp. | | 34 | 37 | 8.03 | |
| Tb927.7.1400 | TAC60 | kDNA segregation | 33 | 61 | 4.95 | TAC |
| Tb927.5.2790 | mt. DNA polymerase beta-PAK | kDNA replication | 32 | 87 | 11.05 | kDNA |
| Tb927.7.5330 | hyp. | | 31 | 83 | 7.74 | cyto. |
| Tb927.8.3840 | hyp. | | 29 | 80 | 6.97 | cell tip + glycos. |
| Tb927.7.3050 | mt. SSU r. prot., put. | translation | 23 | 132 | 6.03 | mito. |
| Tb927.7.6800 | LSU r. prot., mt., put. | translation | 20 | 42 | 8.39 | mito. |
| Tb927.10.15170 | hyp. | mRNA bind. | 20 | 77 | 9.75 | nucleol. |
| Tb927.11.370 | repressor activator prot. 1 | DNA bind. | 20 | 93 | 4.65 | nucl. |
| Tb927.9.7020 | hyp. | | 19 | 25 | 4.93 | mito. |
| Tb927.11.3640 | mt. r. prot. L27 | translation | 19 | 22 | 10.5 | kDNA + mito. |
| Tb927.8.2820 | hyp. | | 18 | 141 | 5.2 | |
| Tb927.10.4080 | hyp. | | 17 | 127 | 7.93 | kDNA + mito. |
| Tb927.10.8890 | kinetoplast DNA-associated prot., put. | DNA bind.; phosphate ion bind. | 17 | 24 | 11.33 | kDNA |
| Tb927.9.4500 | heat shock prot., put. | chaperonne | 16 | 91 | 9.13 | ER |
| Tb927.7.4710 | MRPL46 | translation | 16 | 34 | 8.47 | kDNA + mito. |
| Tb927.11.1250 | mt. SSU r. prot., put. | translation | 15 | 98 | 6.86 | mito. |
| Tb927.10.5300 | eIF-6, put. | ribosome biogenesis | 15 | 27 | 4.8 | nucleol. |
| Tb927.8.6980 | FAZ14 | protein complex bind. | 14 | 95 | 7.46 | FAZ |
| Tb927.11.6000 | MRPL4 | translation | 14 | 53 | 9.2 | mito. |
| Tb927.11.10080 | LSU r. prot., mt., put. | translation | 13 | 22 | 11.92 | mito. |
| Tb927.3.820 | LSU r. prot., mt., put. | translation | 13 | 22 | 10.53 | mito. |
| Tb927.11.14960 | pumilio/PUF RNA bind. prot. 7, put. | RNA bind.; protein bind. | 13 | 79 | 8.56 | nucleol. |
| Tb927.11.8050 | Sas10 C-terminal cont. prot., put. | ribosome biogenesis | 13 | 60 | 5.37 | nucl. |
| Tb927.10.9890 | hyp. | | 12 | 97 | 8.28 | glycos. |
| Tb927.8.4290 | Nop16, put. | ribosome biogenesis | 12 | 22 | 10.97 | nucleol. |
| Tb927.11.6360 | 60S r. prot. L24, put. | struct. constituent of rib. | 12 | 23 | 11.12 | cyto. |
| Tb927.9.6910 | hyp. | | 12 | 18 | 8.67 | kDNA + mito. |
| Tb927.8.1210 | hyp. | | 12 | 78 | 4.57 | |
| Tb927.4.4600 | MRPL43 | translation | 12 | 31 | 10.1 | mito. |
| Tb927.9.15060 | rRNA processing prot., put. | rRNA processing | 11 | 28 | 10.84 | nucleol. |
| Tb927.6.2050 | RRS1, put. | ribosome biogenesis | 11 | 25 | 10.49 | nucleol. |
| Tb927.9.14410 | RNA 3'-terminal phosphate cyclase, put. | mRNA modification | 11 | 39 | 6.75 | nucleol. |
| Tb927.11.2250 | conserved prot., unknown function | mRNA bind. | 11 | 27 | 10.65 | cyto. |
| Tb927.10.13770 | hyp. | | 11 | 13 | 9.78 | kDNA + mito. |
| Tb927.10.9920 | hyp. | | 10 | 44 | 9.73 | nucleol. |
| Tb927.11.11470 | kinetoplast r. PPR-repeat cont. prot. 14 | ribosome biogenesis | 10 | 32 | 7.13 | kDNA |
| Tb927.9.7250 | hyp. | | 10 | 21 | 11.15 | cyto. |
| Tb927.11.13890 | AKAP7, put. | cAMP signaling | 10 | 31 | 10.08 | mito. |
| Tb927.7.870 | hyp. | | 10 | 41 | 7.12 | mito. |
| Tb927.10.14700 | hyp. | mRNA bind. | 9 | 38 | 4.73 | nucl. |
| Tb927.11.10850 | hyp. | | 9 | 48 | 10.68 | kDNA + mito. |
| Tb927.10.600 | mt. r. prot. L29 | translation | 9 | 63 | 7.88 | mito. |
| Tb927.9.10400 | hyp. | mRNA bind. | 9 | 51 | 10.18 | nucl. |
| Tb927.7.320 | RNA-bind. prot. 8 | RNA bind. | 9 | 20 | 9.61 | |
| Tb927.11.11460 | ATOM69 | mito. Import | 9 | 69 | 5.28 | mito. |
| Tb927.10.380 | kinetoplast r. prot. 5 | translation | 9 | 40 | 10.05 | kDNA + mito. |
| Tb927.10.6850 | MRPS18 | translation | 9 | 37 | 10.39 | cyto. |
| Tb927.10.7990 | replication factor C s.u. 3, put. | DNA replication | 8 | 39 | 9.6 | nucl. |
| Tb927.10.4220 | hyp. | telomeric DNA bind. | 8 | 34 | 7.11 | |
| Tb927.7.3960 | MRPL16 | translation | 8 | 19 | 10.75 | kDNA |
| Tb927.11.9450 | peptidyl-prolyl isom., put. | protein folding | 8 | 21 | 6.26 | kDNA + mito. |
| Tb927.7.2940 | histone H2A, put. | DNA bind. | 8 | 14 | 11.89 | nucl. |
| Tb927.4.750 | 50S r. prot. L7Ae, put. | translation | 7 | 16 | 9 | nucl. |
| Tb927.10.7380 | LSU r. prot., mt., put. | translation | 7 | 40 | 10.04 | mito. |
| Tb927.6.3930 | LSU r. prot., mt., put. | translation | 7 | 49 | 9.33 | kDNA + mito. |
| Tb927.10.12050 | LSU r. prot., mt., put. | translation | 7 | 34 | 5.34 | kDNA + mito. |
| Tb927.8.3160 | hyp. | | 7 | 19 | 10.37 | TAC |
| Tb927.4.630 | MICOS complex s.u. MIC34 | cristae formation | 7 | 34 | 6.69 | mito. |
| Tb927.9.7670 | hyp. | | 7 | 26 | 10.42 | nucleol. |
| Tb927.9.4000 | hyp. | telomeric DNA bind. | 7 | 38 | 9.63 | nucl. |
| Tb927.10.13730 | 60S r. prot. L7, put. | translation | 7 | 34 | 10.99 | nucleol. |
| Tb927.11.5500 | mt. SSU r. prot., put. | translation | 7 | 95 | 8.74 | mito. |
| Tb927.11.2190 | MICOS complex s.u. MIC10-2 | cristae formation | 6 | 11 | 9.6 | mito. |
| Tb927.4.470 | snoRNP prot. GAR1, put. | mRNA bind. | 6 | 22 | 12.03 | nucleol. |
| Tb927.7.4140 | MRPL21 | translation | 6 | 21 | 9.42 | kDNA + mito. |
| Tb927.7.1290 | prot. (DUF2012), put. | | 6 | 27 | 6.92 | cyto. |
| Tb927.4.1810 | MRPL33 | translation | 6 | 13 | 11.19 | mito. |
| Tb927.5.3110 | MRPL49 | translation | 6 | 24 | 9.9 | kDNA + mito. |
| Tb927.10.8200 | r. prot. L1p/L10e family, put. | translation | 6 | 31 | 9.44 | cyto. |
| Tb927.8.3750 | Nop56 | snoRNA bind. | 6 | 54 | 8.92 | nucleol. |
| Tb927.10.3030 | proteasome regulatory s.u. 11 | endopeptidase act. | 6 | 34 | 6.52 | cyto. |
| Tb927.8.5490 | Nop52, put. | rRNA processing | 6 | 72 | 9.64 | nucleol. |
| Tb927.10.10590 | histone H2B, put. | DNA bind. | 6 | 13 | 12.29 | nucl. |
| Tb927.9.5150 | r. prot. S6, put. | translation | 6 | 14 | 7.37 | cyto. |
| Tb927.5.4260 | histone H4, put. | DNA bind. | 6 | 11 | 11.65 | nucl. |
| Tb927.10.12330 | zinc finger prot. family member, put. | RNA bind. | 6 | 22 | 9.25 | cyto. |
| Tb927.11.14020 | nuclear RNA bind. domain 2 | RNA / mRNA / protein bind. | 6 | 30 | 11.25 | cyto. |
| Tb927.6.4130 | succinate dehyd. s.u. 3 | ox. phos. | 6 | 12 | 9.02 | mito. |
| Tb927.4.590 | Prot. (DUF1620), put. | | 5 | 88 | 8.96 | ER |
| Tb927.9.5320 | nucleolar RNA bind. prot., put. | mRNA bind.; snoRNA bind. | 5 | 47 | 9.65 | nucleol. |
| Tb927.4.3690 | mt. SSU r. prot., put. | translation | 5 | 49 | 7.43 | kDNA |
| Tb927.2.4980 | EMG1/NEP1 methyltransf., put. | ribosome biogenesis | 5 | 30 | 9.42 | nucleol. |
| Tb927.8.5280 | MRPS34 | translation | 5 | 29 | 5.41 | mito. |
| Tb927.6.2080 | mt. SSU r. prot., put. | translation | 5 | 46 | 9.45 | mito. |
| Tb927.7.4550 | MRPL7/L12 | translation | 5 | 19 | 9.96 | mito. |
| Tb927.10.5620 | fructose-bisphosphate aldolase, glycos. | glycolysis | 5 | 41 | 8.98 | glycos. |
| Tb927.10.7500 | fibrillarin | snRNP biogenesis | 4 | 32 | 10.47 | nucl. |
| Tb927.9.8290 | MRPL30 | translation | 4 | 25 | 10.65 | kDNA + mito. |
| Tb927.7.3430 | peptidyl-prolyl isom., put. | protein folding | 4 | 26 | 6.39 | mito. |
| Tb927.10.11390 | 60S r. prot. L6, put. | translation | 4 | 21 | 11.05 | cyto. |
| Tb927.5.2080 | GMP reductase | GMP reductase act. | 4 | 52 | 9.69 | glycos. + nucleol. |
| Tb927.10.470 | choline dehyd., put. | choline dehyd. act. | 4 | 57 | 7.88 | mito. |
| Tb927.7.7010 | LSU r. prot., mt., put. | translation | 4 | 18 | 11.01 | mito. |

Abbreviations:

AKAP: A-kinase anchor protein; bind.: binding; cAMP: cyclic adenosine monophosphate; cyto.: cytoplasmic; dehyd.: dehydrogenase; DUF: domain of unknown function; eIF: eukaryotic initiation factor; ER: endoplasmic reticulum; FAZ: flagellum attachment zone; glycos.: glycosomal; iso. p.: isoelectric point; isom.: isomerase; methyltransf.: methyltransferase; MICOS: mitochondrial contact site and cristae organisation system; mito.: mitochondrial; MRP: mitochondrial ribosomal protein; mt. SSU: mitochondrial small subunit ribosomal; NOP: nucleolar protein; nucl.: nucleus; nucleol.: nucleolus; prot.: protein; put: putative; r. : ribosomal; RRS1: ribosome biogenesis regulator protein 1; snRNP: small nuclear ribonucleoprotein; struct.: structural; s.u.: subunit; ter.: terminal

**Table S3.** TAC102 yeast two-hybrid screen clones

| Clone Name | Type Seq | Product | Gene Name (Best Match) | Score (A, high confidence, C good confidence) | Chr | Start (nt) | Stop (nt) | Frame | Sense |
|---|---|---|---|---|---|---|---|---|---|
| TB_Ge_hgx5372v1_pB27_A-10 | 5p 3p | | Tb927.9.250 | N/A | 9, C | 645 | -263 | ?? | AntiSense |
| TB_Ge_hgx5372v1_pB27_A-16 | 3p | | Tb927.10.5350 | N/A | 10, C | No Data | 2155 | ?? | AntiSense |
| TB_Ge_hgx5372v1_pB27_A-5 | 5p 3p | NUP109 | Tb927.11.15990 | C | 11, C | 2058 | 3621 | IF | Sense |
| TB_Ge_hgx5372v1_pB27_A-15 | 5p 3p | NUP109 | Tb927.11.15990 | C | 11, C | 2058 | 3621 | IF | Sense |
| TB_Ge_hgx5372v1_pB27_A-2 | 5p 3p | p166 | Tb927.11.3290 | A | 11, W | 111 | 630 | IF | Sense |
| TB_Ge_hgx5372v1_pB27_A-9 | 5p 3p | p166 | Tb927.11.3290 | A | 11, W | 111 | 630 | IF | Sense |
| TB_Ge_hgx5372v1_pB27_A-13 | 5p 3p | p166 | Tb927.11.3290 | A | 11, W | 147 | 2149 | IF | Sense |
| TB_Ge_hgx5372v1_pB27_A-12 | 5p 3p | VSG | Tb927.8.180 | N/A | 8, C | 1409 | 187 | ?? | AntiSense |
| TB_Ge_hgx5372v1_pB27_A-4 | 5p 3p | | Intergene | N/A | 11, W | No Data | No Data | ?? | Sense |
| TB_Ge_hgx5372v1_pB27_A-6 | 5p 3p | | Intergene | N/A | 11, W | No Data | No Data | ?? | Sense |
| TB_Ge_hgx5372v1_pB27_A-17 | 5p 3p | | Intergene | N/A | 11, W | No Data | No Data | ?? | Sense |
| TB_Ge_hgx5372v1_pB27_A-14 | 5p 3p | | Intergene | N/A | 6, W | No Data | No Data | ?? | Sense |
| TB_Ge_hgx5372v1_pB27_A-3 | 5p 3p | | Intergene | N/A | 8, C | No Data | No Data | ?? | Sense |
| TB_Ge_hgx5372v1_pB27_A-8 | 5p 3p | | Intergene | N/A | 8, C | No Data | No Data | ?? | Sense |
| TB_Ge_hgx5372v1_pB27_A-11 | 5p 3p | | Intergene | N/A | 8, C | No Data | No Data | ?? | Sense |

**End section**

**Data, code and materials.** The datasets supporting this article have been uploaded as part of the supplementary material.

**Authors' contributions.** HB carried out the biochemical lab work, participated in the molecular lab work, cell culture, immunofluorescence, design of the study, data analysis and drafted the manuscript; LP participated in the molecular lab work, cell culture and immunofluorescence; TO coordinated the study, participated in data analysis, design of the study and revised the manuscript. All authors gave final approval for publication and agree to be held accountable for the work performed therein.

**Competing interests.** We declare that we have no competing interests.

**Funding.** This work was supported by the Swiss National Science Foundation (17423) and the canton of Bern.

**Acknowledgments.** We thank Roman Trikin and Ana Kalichava for technical assistance, Simona Amodeo and Irina Bregy for reading the manuscript. We thank Andre Schneider and Volker Heussler for antibodies. We also thank the Proteomic and Mass Spectrometry Core Facilities (PMSCF) and the Microscopy Imaging Centre (MIC) from the University of Bern, Switzerland.
